# Unmasking Early Microglial Remodeling in an Alzheimer’s Disease Mouse Model

**DOI:** 10.1101/2025.10.07.681006

**Authors:** Priyanka Saminathan, Sara McArdle, Maija Corey, Namratha Nadig, Camille Fang, Alicia Gibbons, Mahati Rayadurgam, Sonia Sharma

**Affiliations:** La Jolla Institute for Immunology, La Jolla, CA 92037, United States of America

**Author notes:** **Correspondence:** Sonia Sharma, Sara McArdle.

**Keywords:** Microglia morphometry, Image segmentation, QuPath, Neuroinflammation, Alzheimer’s Disease, Iba1 and CD68 quantification, Morphological remodeling, Hippocampus

## Abstract

Early neuroimmune remodeling is a critical yet understudied component of Alzheimer’s disease (AD) pathogenesis. To investigate microglial contributions to AD development prior to overt plaque deposition, we developed an open-source morphometric pipeline to systematically quantify hippocampal microglial structure and activation states in pre-plaque 5xFAD mice. Across ∼11,000 cells, we extracted multidimensional parameters including area, circularity, convex hull, branch points, nearest-neighbor distance, and nuclear features, alongside Iba1 and CD68 intensity measurements. While no significant overt gliosis was observed at this early stage, microglia from 5xFAD mice exhibited subtle trends toward increased structural complexity compared to wild-type controls. Importantly, significant sex-specific differences were detected within the CA1 subregion: male 5xFAD microglia displayed hyper-ramified morphologies consistent with enhanced surveillance states, whereas female microglia demonstrated greater density and a more reactive phenotype. Correlation analyses revealed a conserved association between microglial complexity and Iba1/CD68 expression, independent of sex or genotype, underscoring a fundamental link between cytoskeletal remodeling and phagolysosomal activity. These findings highlight the capacity of morphometric profiling to sensitively detect early, region-specific, and sex-dependent shifts in microglial phenotype before amyloid deposition. By integrating quantitative morphology with canonical molecular markers, this framework provides a robust and unbiased approach for characterizing microglial activation trajectories. Such early readouts may inform biomarker discovery and therapeutic strategies aimed at modulating microglial responses to delay or prevent AD progression.

## 1 Introduction

Neuroinflammation is a core dimension of Alzheimer’s disease (AD) pathobiology, with microglia, the brain’s resident macrophages, poised at the interface of risk, injury detection, and circuit remodeling^1^. Genetic studies first implicated microglia in Alzheimer’s disease, as many AD risk variants map to microglial pathways. Notably, rare loss-of-function mutations in TREM2 markedly increase susceptibility and hasten disease progression, alongside additional variants affecting immune regulation and lipid-handling genes that converge on microglial function.(e.g., CD33, CR1, ABCA7). These data argue that microglial signaling pathways can shift the disease trajectory rather than simply report downstream damage^2–7^. Functionally, early microglial responses shape the sequence from amyloid-β (Aβ) stress to synaptic failure and tauopathy. In mouse models, complement components such as C1q and C3 accumulate at synapses before overt plaques^8,9^. Microglial complement signaling then drives early synapse elimination and memory deficits, while blockade of complement protects synapses and neurocognitive behavior^8,10^. These studies situate microglia as proximal orchestrators of the earliest circuit changes that better correlate with cognition status than bulk amyloid burden. Microglia also facilitate tau spread via exosome release, while microglial ablation or inhibition of exosome biogenesis curtails propagation *in vivo* ^11,12^, further strengthening the link between innate immunity to proteopathic spread in AD.

At the cellular level, microglial state transitions are accompanied by stereotyped structural remodeling that encodes functional roles in surveillance, synapse interaction, and phagocytosis. During homeostasis, microglia display fine, highly ramified arbors with small somata and dynamic process motility; stressors and disease cues drive shifts through hyper-ramified, deramified/bushy, and amoeboid phenotypes^13–15^. At early, preplaque stages in AD, *in vivo* imaging shows microglia adopting these hyper-ramified, primed morphologies with heightened process motility and clustering around emerging Aβ seeds^16–19^. Microglia accelerate rapid plaque compaction and form barrier-like assemblies that limit amyloid spread and neuritic dystrophy, suggesting a context dependent balance between containment and collateral damage^18,20,21^. Understanding when and where this balance tips remains a central challenge for early-stage intervention in AD. Translationally, the field requires biomarkers and analytic frameworks that detect subtle, pre-plaque immune remodeling with spatial precision, quantify cell-state heterogeneity, and link these readouts to synaptic health and future lesion emergence.

High-throughput imaging pipelines for microglial morphology, such as the Convolutional Neural Network (CNN)-based classifiers developed in ischemia, demonstrate scalable, unbiased labeling and correctly capture hallmark activation transition^22^. However, fixed morphotype categories and non-AD validation can obscure the subtle, spatially patterned, and sex-dependent remodeling that defines early AD^17,18^. Accordingly, resolving early AD-relevant shifts requires continuous, feature-level morphometrics coupled to Iba1/CD68 and spatial statistics.

A second, underexplored dimension of microglial modulation is biological sex. Microglia exhibit sex-dependent developmental trajectories and transcriptomes even at baseline; males and females differ in microglial density, morphology, and immune reactivity across brain regions^23–25^. In parallel, APOE-ε4 confers a greater AD risk and stronger association with tauopathy in women, suggesting interactions between sex, lipid biology, and innate immune tone that could bias disease initiation and spread^26–29^. Notably, female 5xFAD mice develop plaques earlier and show more pronounced gliosis, underscoring that sex differences are evident even in experimental models^30,31^. These observations motivate sex-aware analyses of early microglial remodeling in AD models and human cohorts.

Building on this framework, we focus on early microglial remodeling as a predictor and potential driver of downstream AD pathology using the 5xFAD mouse model. 5xFAD mice overexpress human APP (Swedish, Florida, London) and PSEN1 (M146L, L286V), and begin depositing extracellular Aβ by 2 months, with robust early microgliosis^32^. Despite lacking tau pathology and progressing faster than human disease, their aggressive amyloidosis and early microglial activation make them a useful system for probing pre-plaque and early-stage remodeling^33^. We hypothesize that in pre-plaque stages, microglia undergo region-specific and sex-modulated structural transitions, characterized by hyper-ramification in vulnerable circuits and local changes in cell density and spacing. We believe these findings foreshadow complement-tagged synapse loss and later Aβ pathology. To test this, we deploy a quantitative, open, and reproducible immunofluorescence image analysis pipeline optimized for the study of microglia at ages preceding dense plaque deposition. Our image-analysis workflow combines semi-automated segmentation with per cell morphometrics which include : cell and soma area/circularity, convex hull area/solidity, total process length, branching index, critical radius and value (Sholl analysis), Schoenen Ramification Index (SRI), cellular Iba1 and CD68 intensities, and population-level spatial statistics that include microglial density and nearest-neighbor distance (NND). Methodologically, this study leverages validated tools to enhance sensitivity to changes that are typically lost in 2D sections. Automated or semi-automated platforms reduce observer bias and scale to thousands of cells, while recent benchmarking underscores the discriminative value of convex hull and skeleton based features and encourages open, machine-learning-ready data standards. These advances, coupled with region resolved sampling, allow us to map the topography of microglial remodeling across vulnerable circuits with statistical power.

## 2 Methods

### 2.1 Animal handling

Transgenic 5xFAD SJL mice were purchased from Jackson Laboratories. Mouse experiments were performed with 6-8 week old mice of both sexes housed in specific-pathogen-free conditions at the La Jolla Institute for Immunology. All procedures involving mice were performed according to the Institutional Animal Care and Use Committee (IACUC) at La Jolla Institute for Immunology.

### 2.2 Histology

Mice were transcardially perfused with 10 ml PBS and 10 ml 4% PFA, and brains were removed from the skull and fixed for additional 48 hours. Following washing with PBS, brains were cryoprotected in 30% sucrose solution in PBS and frozen in OCT compound. Coronal sections were cut at 35 μm and stored in cryoprotectant (30% PEG300, 30% glycerol, 20% 0.1 M phosphate buffer, and 20% ddH2O) at -20C until staining. Sections were washed with PBS (2x10 min), and incubated with 10% normal donkey serum with 0.5% Tween 20 in PBS for 2h at RT. This and subsequent incubations were carried out with a laboratory rocker. Sections were transferred to a 2 ml tube for incubation with the primary antibody cocktail (rat anti-CD68, clone FA-11, 2 µg/ml, Biolegend cat #137001, and rabbit anti-Iba1, clone E404W, 0.2 µg/ml, Cell Signaling Technologies cat#17198S) overnight at 4C. Sections were washed with PBS with 0.1% Tween 20 (PBS-T, 3x5 min) using transwell membrane inserts to facilitate transfer of sections to new wells of a multiwell plate. Samples were stained with goat anti-rat IgG Alexa Fluor 568, and goat anti-rabbit IgG Alexa Fluor 647 (both at 4 µg/ml, ThermoFisher Scientific cat# A11077, and A32733, respectively) cross-adsorbed secondary antibodies for 2h at RT. Samples were washed with PBS-T (2x5 min), PBS (1x5 min), stained with Hoechst 33258 (10 µg/ml, 10 min), washed with PBS (3x5 min), and embedded in Prolong Gold using #1.5 coverslip. Images were captured on a ZEISS Axioscan 7 slides scanner with a 20x 0.8NA objective, using 385, 567, 630 nm LED illumination modules, a multiband filter cube, and a 712 M camera, resulting in Z-stacks with a 0.172 x 0.172 x 0.490 µm voxel size.

### 2.3 Image pre-processing

To reduce noise, improve spatial resolution, and reduce the effect of out-of-focus light, Z-stacks were deconvolved using the Huygen’s Essential software suite (v23.10, Scientific Volume Imaging). First, individual tiles or each channel were deconvolved with the quick maximum likelihood estimate algorithm, using a brick mode of 1. A maximum intensity Z-projection was calculated for each tile and saved as an .ome.tiff. The 2D projections were stitched together using a single slice from the original .czi image as a starting template. The final file was saved as a 32-bit, 3 channel .ome.tiff image.

### 2.4 Microglia segmentation

All image analysis was performed in QuPath v0.5.1^34^. The hippocampus region was manually annotated, excluding the granule cell nuclei layer in the dentate gyrus. The CA1 subregion was also outlined. Within the full hippocampus region, microglia segmentation proceeded in 2 steps. First, a pixel classifier was trained on the Iba1 channel to segment microglia processes as precisely as possible and detection objects were created (minimum size threshold = 0.2 μm). Separately, nuclei were segmented with Cellpose^35^ using the ‘nuclei’ model on the Hoechst channel through the QuPath Cellpose extension.^36^ Microglial nuclei were identified as those that were at least 30% Iba1 positive by area, as measured with the Iba1 pixel classifier; all other nuclei were deleted.

A custom script merged the separate Iba1 and nucleus detections into microglia (see Github: https://github.com/saramcardle/Image-Analysis-Scripts/tree/master/Microglia%20Analysis%20in%20QuPath). Briefly, we collected all detections that were within 100 μm of a microglia nucleus. Nearby Iba1 detections were merged using binary dilation and contraction to fill in small gaps that may have arisen from incomplete labeling of thin dendrites. The area that was contiguous with the nucleus was segmented, labeled as a microglia, and expanded to cover the entire nucleus. Looping over all Iba1+ nuclei resulted in a set of overlapping microglia objects; any remaining Iba1+ objects that were not associated with a nucleus were deleted. Overlapping microglia were separated with watershed separation using the nuclei centroids as initial markers. Each nucleus and its associated microglia outline were assigned matching names. Finally, to reduce artifacts, cells were removed if the nucleus was touching the hippocampal boundary or if the nuclear area or total cell area was too small (<5 μm^2^ or <50 μm^2^, respectively), suggesting it was a cell fragment.

### 2.5 Microglia metrics

Built-in QuPath functions were used to measure the area, perimeter, major and minor Feret diameters, and convex hull area and perimeter of each cell and nucleus, from which we calculated circularity, solidity, convexity, and aspect ratio (see ^22^ for definitions). The area of each cell that was positive for CD68 (pixel intensity >1000 AU) and the average Iba1 fluorescence intensity were calculated. More complex cell features were measured in Fiji.^37^ Each cell was exported from QuPath as a binary image with the nucleus saved as an overlay. In Fiji, the soma region (nucleus expanded by 3 pixels) was deleted, the remaining processes were skeletonized, and the skeleton was analyzed^38^ for length and branch points. Additionally, the Simple Neurite Tracer plugin^39^ was used to calculate Sholl features^40^ of the microglia shape, using the nucleus centroid as a starting point and measuring concentric circles every 0.25 μm from the edge of the nucleus out to 50 μm. All metrics were calculated based on the sampled data and Schoenen Ramification Index was defined as the critical value divided by the number of branches. Branch thickness was calculated as the cell cytoplasm area (cell area minus nuclear area), divided by the total branch length and the nuclear offset was calculated as the distance between the cell centroid and nucleus centroid.

Spatial relationships between cells were calculated in QuPath v0.5.1 using Delaunay triangulation. The mean neighbor distances and mean triangle areas were calculated for the cell centroids, using a maximum distance of 250 μm. Separately, to identify clumped cells, Delaunay triangulation was calculated for the nucleus objects themselves, with a maximum distance of 25 μm. Reactive microglia were identified as those with CD68 expression >=35% area, a Branching Index >=1, and a nucleus <25um from at least one other nucleus.

### 2.6 Statistical Analysis

The shape, intensity, and spatial distribution metrics for all cells were exported as .csv files, which were loaded into Matlab (Mathworks) for further processing. In most cases, each hippocampus (or CA1 region) was summarized as a median of all cells in that region. The statistical significance of group comparisons were calculated in Prism v10 (GraphPad) using either a 2-way ANOVA of the slide medians, with uncorrected Fisher’s LSD with single pooled variance .

To look at the relationships between parameters, the dataset of 10490 microglia were divided into deciles of 1049 cells based on area, circularity, or branching index. The cells in each decile group were split by genotype and then the average CD68% or Iba1 intensity of the cells in each group were calculated. To evaluate the profile of the reactive microglia, the 50 detected reactive cells were compared to the homeostatic cells from the CA1 region of the 3 matching female 5xFAD mice that had a significant number of reactive cells. Outliers were removed using the ROUT test (with Q=1%) and then shape metrics were compared with an unpaired, two-tailed t-test.

## 3 Results

### 3.1 Workflow for assessing structural changes in microglial activation

Briefly, 35 um sections from frozen mouse brains were cryosectioned and immunostained for CD68 and Iba1 using a free-floating method (Fig 1A). They were imaged with a widefield slidescanner, acquiring Z stacks with a 0.49 µm step size. The Z stacks were deconvolved and projected (Huygen’s Essential, Scientific Volume Imaging) to form a single image that captured a large depth of field. The resulting image contained more microglial processes than a single focal plane, but with improved clarity and signal-to-noise ratio compared to a maximum intensity projection of the entire stack (Fig. 1B) Segmentation and quantification of the microglia loosely followed the process in previous work^22^, but used the open-source analysis platform QuPath^34^ for easier reproduction and adaptability. The code is available at Github. Microglia segmentation proceeded in 2 steps (Fig 1C). First, nuclei were segmented with Cellpose^35^ and filtered for Iba1 expression. Simultaneously, a pixel classifier was used to find all Iba1+ processes. Neighboring objects were merged through morphological closing, cells were filtered for those associated with an Iba1+ nuclei and overlapping cells were separated. The cell intensities, shape features, and spatial relationships were measured in QuPath and Fiji. A summary of microglial morphological and spatial parameters, and their association with microglial activation states, is shown in Supplemental Table 1.

**Figure 1.**
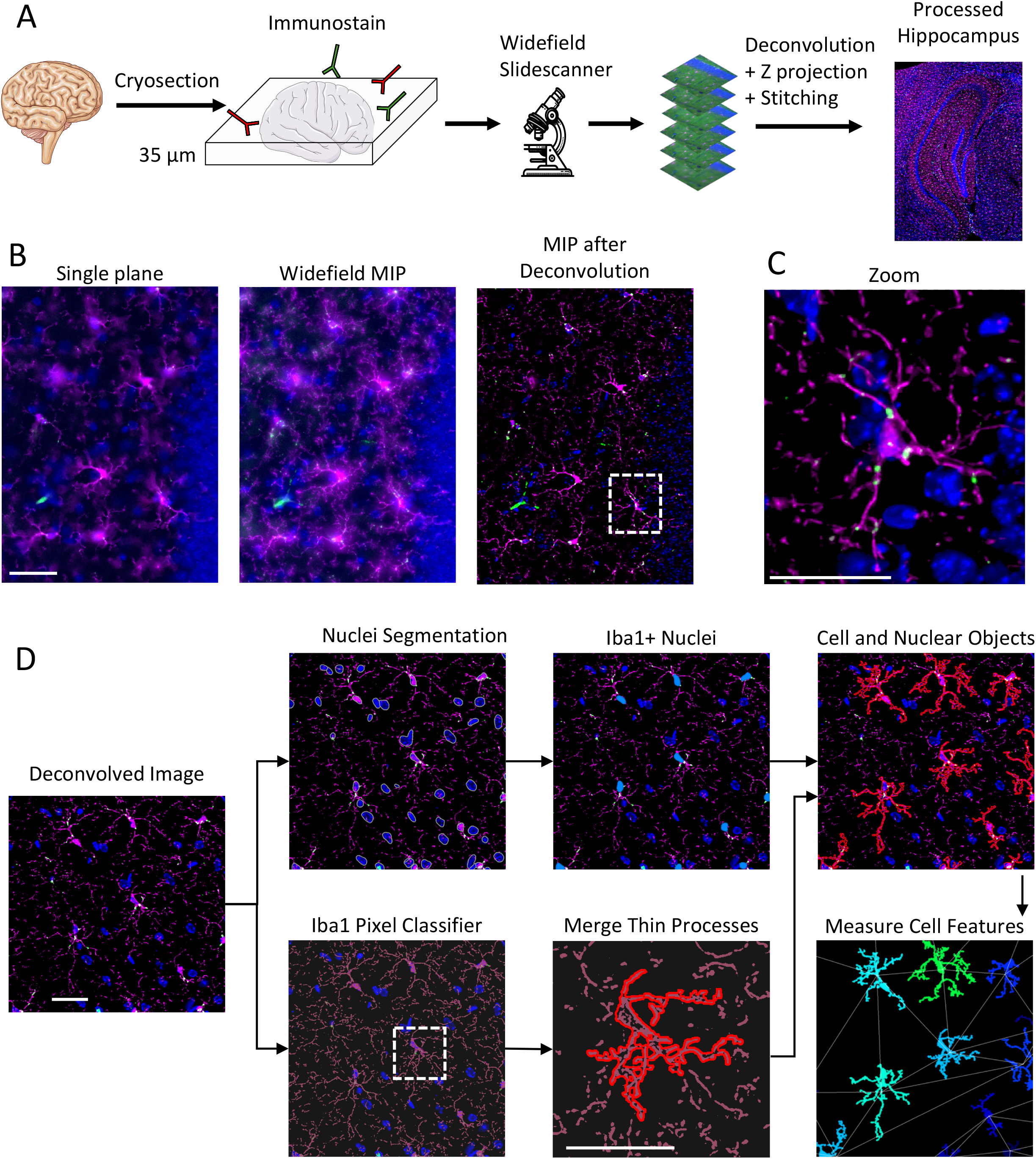
Workflow for assessing structural changes in microglial activation. A) Free-floating 35-µm cryosections were immunostained for Iba1 and CD68 and imaged on a widefield slide scanner as Z-stacks (Δz = 0.49 µm). Tiles were deconvolved, maximum-intensity projected, and stitched to generate extended-depth, whole-hippocampus images. Scale bar = 50 µm. Icons come from biorender.com. B) Comparing single focal plane, maximum-intensity projection, or deconvolved maximum-intensity projection images shows that the deconvolution process yields higher spatial resolution and better signal-to-noise than the single plane or projected widefield images. Hoechst = blue, Iba1 = magenta, CD68 = green. Look-up tables are identical for the first two images, but adjusted for the deconvolved image to prevent oversaturation. Scale bar = 25 µm. C) A zoom-in on the marked region of the deconvolved image shows crisp cell shape details and CD68 localization. Scale bar = 25 µm. D) Microglia segmentation used a two-step pipeline: nuclei were segmented with Cellpose while a trained Iba1 pixel classifier delineated microglial arbors. Morphological filters merged thin processes and split touching objects. The cell outlines and nuclei were combined through the entire hippocampus and cell intensity features and shapes were measured, along with spatial relationships.

### 3.2 Early hippocampal microglia remodeling in 5xFAD Mice: Trending toward hyper-ramification in male 5XFAD microglia and increased density in females

The average hippocampus Iba1 and CD68 intensity gives an overall measure of microglial content. There were no significant differences between genotypes or sexes in either marker in the full hippocampal region, though the Ctrl females had slightly more Iba1 expression than the Ctrl males (Fig 2A). Focusing on segmented microglia, there were no statistically significant differences in average Iba1 or CD68 expression per microglia between sexes or genotypes (Fig 2B). However, the density of segmented microglia was slightly larger in the 5xFAD mice, though this is not seen in the Delaunay mean neighbor distance (Supplementary Fig 1A).

**Figure 2.**
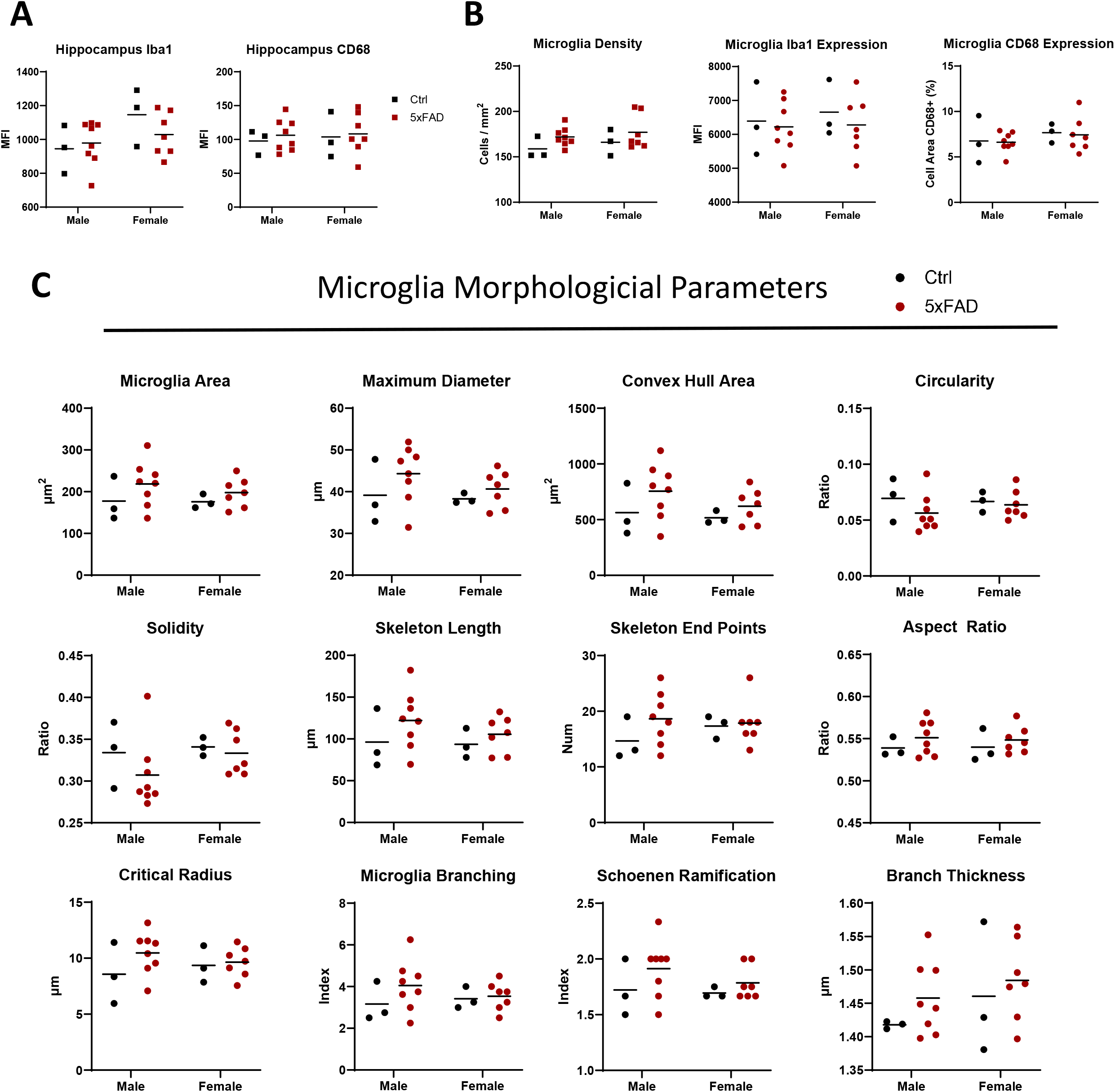
Early hippocampal microglia remodeling in 5xFAD Mice: Trending toward hyper-ramification in male 5XFAD microglia and increased density in females. A) Iba1 and CD68 immunoreactivity were quantified across the whole hippocampus to estimate total hippocampal microglial content and activation. Microglia were segmented and B) the Iba1 Intensity and CD68 positive area, and C) morphological features were measured per cell. Each square represents a measurement of 1 mouse; each circle represents the median value of all microglia in 1 mouse. * = p<.05, ** = p<.01 by 2-way ANOVA.

Looking beyond fluorescence intensity towards shape descriptors, we see no statistically significant differences between groups, but consistent trends towards hyper-ramification in the 5xFAD mice (Fig 2C). Microglia from 5xFAD brains are larger (higher cell area, diameter, convex hull area), more complex (lower circularity and solidity, higher skeleton length and number of skeleton end points) and more branched (higher aspect ratio, critical radius, branching index, and Schoenen Ramification Index), but with thicker branches, all features known to be correlated with hyper-ramified microglia^41^. Furthermore, the 5xFAD males, but not females, had a larger distance between the microglia centroid and the nuclear centroid, suggesting asymmetric branch growth (Supplemental Fig 1B).

To understand how traditional microglia marker expression correlates with shape descriptors, the total population of microglia in the dataset (10,490 cells) were divided into deciles based on area, circularity, and branching index. The average CD68-positive area and average Iba1 intensity were calculated for the cells in each decile for each genotype. In both groups, CD68 and Iba1 expression correlate with decreasing area, increasing circularity, and decreasing branching index (Supplemental Fig1C). Thus, microglial morphometrics trend with Iba1 and CD68 intensity, independent of sex and genotype.

### 3.3 Sex-specific changes observed in the CA1 hippocampal region

Hippocampal atrophy is an early, robust imaging biomarker of AD that correlates with episodic memory decline and clinical progression. Although oligomerization and fibrillization of β-amyloid (Aβ) most often begin in association neocortex, Aβ and tau pathology subsequently involve medial temporal circuits, with early tau spread across entorhinal to hippocampal subfields including CA1^42,43^. CA1 pyramidal neurons constitute a principal hippocampal output stage within the trisynaptic circuit, supporting temporal coding, sequence memory, and spatial navigation along with high metabolic demand and dense glutamatergic/NMDA signaling, contributing to selective CA1 vulnerability^42,43^. Human and animal studies consistently show early CA1 synaptic/neuronal loss and neurofibrillary tangle accumulation that track with memory impairment^42,43^. Microglia are key effectors of this vulnerability in CA1. In 3xTg-AD and other models, microglial density and activation rise in CA1 around the time plaques and tangles emerge, with CA1 showing stronger glial reactivity and neurodegeneration than neighboring subfields^42,43^. Mechanistically, complement-tagged synapses in the hippocampus are aberrantly engulfed by microglia early in disease, providing a causal route from microglial activation to CA1 synapse loss and cognitive decline. These studies show that the CA1 region is central to hippocampal computation, displays early and selective degeneration in AD, and that microglia-mediated synaptic pathology is tightly linked to memory loss^42,43^.

Looking at morphological changes in the microglia in the CA1 region, we see many of the trends for more complex shapes in the 5xFAD mice are more pronounced than in the full hippocampus, particularly in male mice. The overall microglia density was highest in the female 5xFAD mice (not statistically significant, Fig 3A), correlating with the lowest mean Delaunay distance (Supplemental Fig 1D), though there were no significant changes observable either in the cellular Iba1 or CD68 expression (Fig 3B). Male 5xFAD microglia had significantly lower circularity and solidity metrics and higher branching index when compared to the male Ctrl microglia (p<.05 for all, Fig 3C). Male mice also showed a larger distance between the nucleus and full cell centroid (Supplemental Fig 1E). Furthermore, many of the parameters associated with hyperramification of 5xFAD mice are more pronounced in the CA1 region than in the full hippocampus, including cell size, aspect ratio, critical radius, SRI, and branch thickness (Fig 3D).

**Figure 3.**
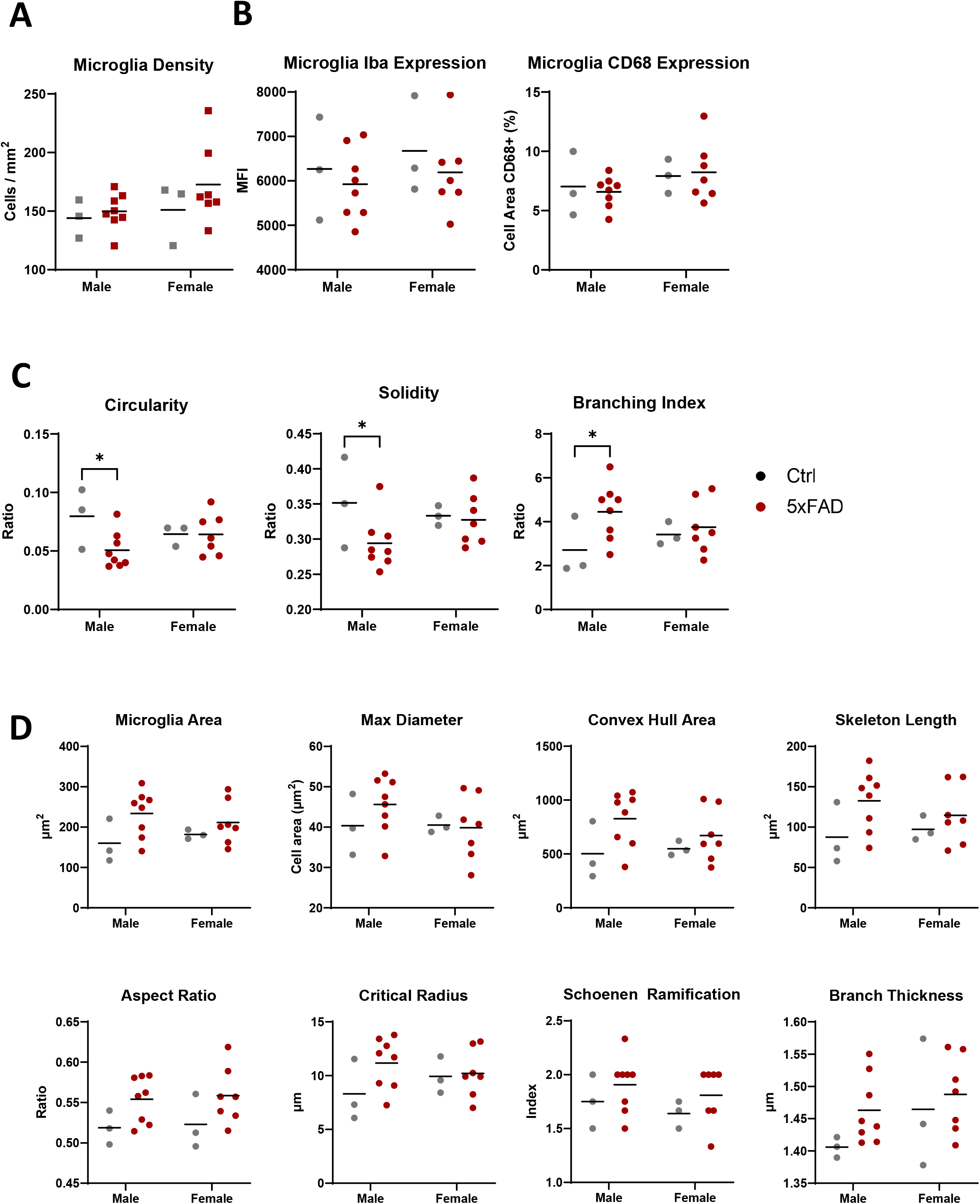
Sex-specific changes observed in the CA1 hippocampal region. A) The overall density of microglia in the hippocampal CA1 region of male and female, 5xFAD and Ctrl mice. B) Microglia from the CA1 region of male 5xFAD mice show significant changes in solidity, circularity, and branching index compared to Ctrl mice. C) Other morphological parameters show a trend towards 5xFAD microglia being more hyperramified than controls, with a larger difference in males. Each square represents a measurement of 1 mouse; each circle represents the median value of all microglia in 1 mouse. * = p<.05, ** = p<.01 by 2-way ANOVA.

### 3.4 Defining early reactive microglia in the female 5xFAD mice

The emergence of highly reactive, amoeboid microglia, typically localized around amyloid plaques during later stages of pathology in AD mouse models is one of the hallmarks of disease progression. Studies demonstrate that by 2–4 months of age, coinciding with the initial stages of plaque formation, microglia in 5xFAD mice adopt an amoeboid morphology characterized by retracted, thickened processes and increased immunoreactivity for activation markers such as Iba1 and F4/80, consistent with a reactive state^8,17,18,44,45^. Furthermore, microglial transcriptome changes in 5xFAD mice by 4–6 months indicate the upregulation of inflammatory and immune genes, complement and integrin signaling, and genes linked to phagocytosis. This is consistent with reactive/disease-associated microglia phenotypes^8,17,18,44,45^. In our current study of 6-8 week old mice, we noticed infrequent cells that have high CD68 expression (>=35% positive), at least medium branching (branching index >=1), and that are clustered together (nucleus centroid <25µm from another nucleus centroid) that appear to be active/reactive microglia (Fig 4A). The female 5xFAD mice showed a greater incidence rate of reactive microglia in the CA1 region than in the rest of the hippocampus or compared to male 5xFAD mice or Ctrl mice (Fig 4B). In the 3 female 5xFAD mice with a substantial frequency of these reactive cells, we found 50 cells across the CA1 regions. Comparing the feature profile of this subset of microglia against the remaining microglia in these 3 mice (CA1 region only), we can see that these cells have the typical reactive microglia profile: higher Iba1 expression, slightly contracted with smaller area, hull area, skeleton length, and critical radius but higher circularity, solidity, and branch thickness. These cells were found almost exclusively in the outer section of the CA1 region (Fig 4D). Thus, although we did not find classically amoeboid microglia in 5xFAD mice at this stage in AD development, we did detect a shift toward early reactivity, specifically in the female 5xFAD CA1 region. These findings support our hypothesis (grounded in prior reports of earlier and stronger microglial activation in female AD models) that female microglia transition to reactive phenotypes sooner than males.

**Figure 4.**
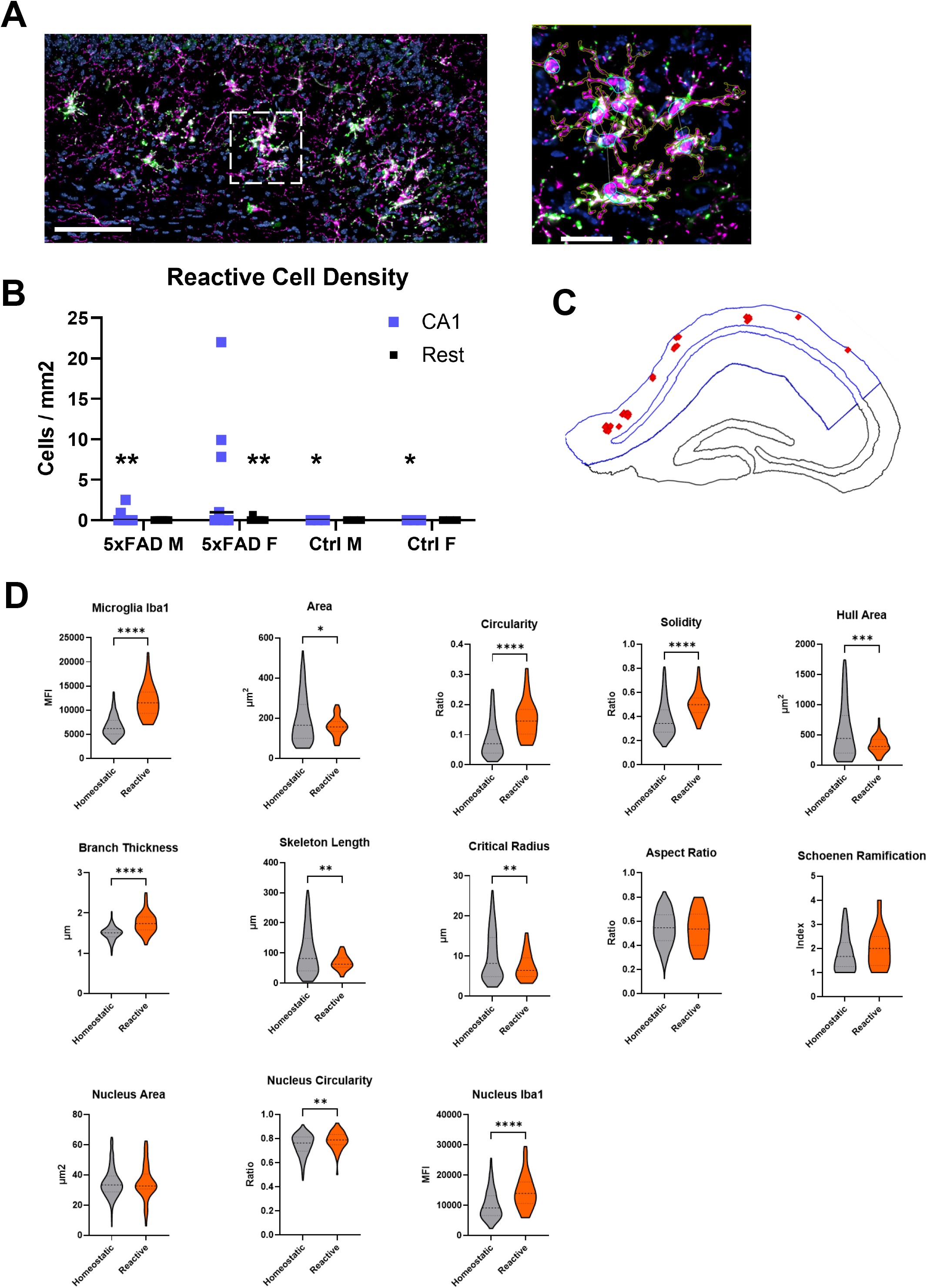
Defining early reactive microglia in the female 5xFAD mice. A) (Left) An image from the CA1 region of a female 5xFAD mouse showing reactive microglia-with visibly high CD68, thick branches, and clustered nuclei. Scale bar = 50 µm. (Right) A zoom-in on the marked region, with the cell and nuclear segmentation visible (brown and cyan, respectively). Scale bar = 25 µm. Hoechst = blue, Iba1 = magenta, CD68 = green. B) The density of all reactive microglia, separated by location, genotype, and sex. Asterisks show significant differences compared to the female, 5xFAD CA1 region. * = p<.05, ** = p<.01 by 2-way ANOVA. 3 of the mice have substantially more reactive microglia than the others. Comparing the morphological and intensity features of the 50 reactive cells found in these mice against the remaining microglia in these 3 mice (CA1 region only), we can see that these cells have changes typical of reactive microglia compared to homeostatic: thicker branches, smaller size, more circular shape, and higher Iba1 expression.

## 4 Discussion

In this study, we provide evidence that microglial remodeling at the pre-plaque stage of 5xFAD mice is both measurable and sex-dependent, underscoring the value of our open-source morphometric pipeline. We observed that male 5xFAD mice exhibited hyper-ramified microglia, whereas female 5xFAD mice displayed a more reactive phenotype, particularly in the CA1 region. These findings align with prior literature demonstrating that microglial changes emerge well before overt amyloid plaque deposition. Boza-Serrano et al. reported that inflammatory alterations are detectable in microglia of 5xFAD mice as early as six weeks, preceding amyloid accumulation^46^. Similarly, Chithanathan et al. documented early microglial activation and morphological changes in two-month-old male 5xFAD mice^47^. Our finding of hyper-ramification in males may therefore represent an exaggerated surveillance response to a subtly changing neuroenvironment, consistent with a primed yet not fully reactive microglial state. In contrast, female 5xFAD microglia more closely resemble disease-associated microglia. Disease-associated microglia have been characterized by increased glycolytic metabolism and antigen presentation, while having reduced phagocytic activity^48^. This sex-biased divergence is consistent with clinical and preclinical observations that females often demonstrate heightened neuroinflammation and accelerated amyloid pathology^49^. By capturing these structural differences with quantitative parameters, our tool allows us to dissect early microglial transitions in a manner that is reproducible and scalable across datasets. Importantly, correlations between microglial structural complexity and canonical activation markers, including Iba1 and CD68, are maintained irrespective of sex or genotype (Supplemental Figure 1C). This conserved association underscores a fundamental coupling between cytoskeletal remodeling and phagolysosomal activity, reinforcing morphological features as quantifiable indicators of microglial functional state. Collectively, these data establish morphometric profiling as a sensitive and unbiased approach to detect early microglial activation, while capturing sex-specific trajectories before amyloid deposition.

In the present study, we focused on developing an image analysis method to semi-automatically quantify microglial parameters and activation states in murine hippocampuses. We characterized >10,000 microglia in 21 mice, making this among the largest studies of microglial shape to date^50^. Performing the analysis on the entire hippocampus, instead of individual ROIs, allowed us to find rare reactive cells in young mice. By making the code available open-source, we hope to provide a useful tool to the community. Due to the low number of reactive cells present in the dataset, we could not rely on a supervised or unsupervised classifier to identify cell phenotypes, as was done in previous work^22,41^. Instead, we used simple thresholds on only a few parameters that can easily be repeated on smaller datasets. Frequently, microglia analysis is performed on single focal planes (getting only a fraction of each cell)^22,41^, widefield Z projections (which blurs fine details), or confocal Z stacks (low-throughput)^22,41^. By acquiring a large Z range, and using deconvolution to reduce the effects of out-of-focus light, we were able to get more of the microglia branching at the speed of widefield imaging. While the software we used for deconvolution in this work is commercial, similar processing can be implemented in open-source software^51^. Here, we also introduce two uncommon metrics for microglia: centroid offset (the distance between the cell centroid and nucleus centroid) and Delaunay mean distance (a measure of cell density). We note that microglia in the CA1 region of 5xFAD mice show higher centroid offset while also trending towards hyperramification. Similarly, reactive cells in the female 5xFAD mice tend to be more clustered. Future work will be needed to determine if this pattern is coincidental or consistent.

Although we did not directly investigate lipid alterations, this analytical framework provides a foundation for future studies aimed at integrating peripheral bioactive lipid profiles with microglial remodeling in murine AD models. Prior work has demonstrated that microglia are key contributors to lipid-mediator signaling: for example, microglial production of leukotrienes has been shown to exacerbate neuroinflammation in APP/PS1 mice, and genetic or pharmacological inhibition of 5-Lipoxygenase and its activating protein reduces pathology and improves outcomes^52^. Conversely, specialized pro-resolving mediators derived from polyunsaturated fatty acids are diminished in AD and, when supplemented, can restore microglial phagocytosis and synaptic function^53^. These findings highlight the importance of lipid-microglia interactions as both biomarkers and therapeutic entry points. By applying our morphometric tool in conjunction with serum lipidomics, we aim to identify correlations between distinct microglial activation states and circulating lipid mediators. Such integration has the potential to define early molecular signatures of disease, inform patient stratification, and uncover druggable pathways that shift microglia from detrimental to homeostatic states. In this way, our current work not only advances methodology for microglial characterization but also lays the groundwork for mechanistic and translational studies linking neuroinflammation, lipid metabolism, and therapeutic targeting in AD.

## 5 Data Availability Statement

The dataset generated for this study can be found in the BioImage Archive at S-BIAD2332. The custom script that merged the separate Iba1 and nucleus detections into microglia is available at Github: https://github.com/saramcardle/Image-Analysis-Scripts/tree/master/Microglia%20Analysis%20in%20QuPath

## 6 Ethics Statement

All animal procedures were approved by the La Jolla Institute for Immunology Institutional Animal Care and Use Committee (IACUC) and conducted in accordance with the National Institutes of Health guidelines for the care and use of laboratory animals. All efforts were made to minimize animal suffering and to use the minimum number of animals required for valid statistical analysis.

## 7 Author Contributions

PS and SS conceptualized and designed the study. PS and SM developed the methodologies. PS, NN and CF conducted the mouse experiments. SM designed image analysis methods and wrote the novel code. PS, SM and SS were responsible for data interpretation and data curation. The original draft of the manuscript was written by PS and SM. SS, MC, AG, and MR contributed to reviewing and editing the manuscript. Funding acquisition and overall supervision of the project were provided by SS. All authors reviewed, edited, and approved the final manuscript.

## 12 Acknowledgments

We would like to thank Kasia Dobaczewska for expert assistance in slide preparation and Dr. Zbigniew Mikulski for image acquisition.

## Figure legends

**Supplemental Figure 1**

A) Delaunay clustering was performed on all microglia in the hippocampus based on the cell centroids. For each cell, the mean distance to its neighbors was calculated in QuPath. Each dot shows the median value of all cells in 1 mouse. B) The distance between the cell centroid and the nucleus centroid was also calculated for each cell. C) By dividing the microglia into deciles based on area, circularity, and branching index, we can see a clear trend in these morphological parameters with CD68 and Iba1 expression. D and E) Delaunay clustering and nuclear centroid offset was calculated on the cells in the CA1 region alone.

**Supplementary Table 1**

Summary of parameters used to assess microglial morphology, complexity, and branching intensity. Cell area, diameter, and convex hull area constitute microglial size parameters. Circularity, solidity, skeleton length, and number of skeleton end points constitute complexity parameters. Aspect ratio, critical radius, branching index, and SRI constitute branching parameters. References supporting parameter descriptions are provided.

