## Supplemental Data for "Unmasking Early Microglial Remodeling in an Alzheimer’s Disease Mouse Model"

*Supplementary Material*

### 1.1 Supplementary Figure 1

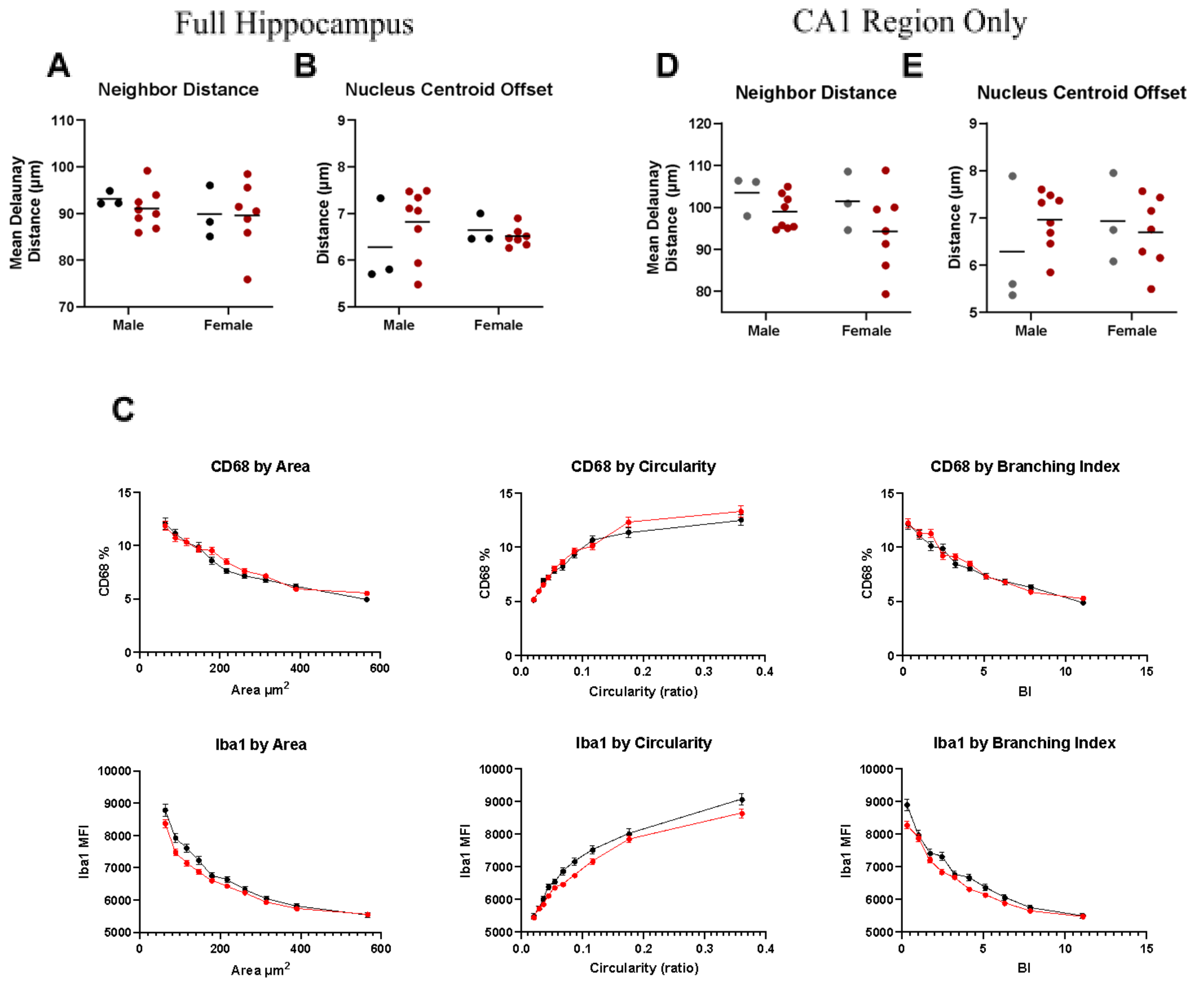


**Supplementary Figure 1**
Delaunay clustering was performed on all microglia in the hippocampus based on the cell centroids. For each cell, the mean distance to its neighbors was calculated in Qu Path. Each dot shows the median value of all cells in 1 mouse. B) The distance between the cell centroid and the nucleus centroid was also calculated for each cell. C) By dividing the microglia into deciles based on area, circularity, and branching index, we can see a clear trend in these morphological parameters with CD68 and Iba1 expression. D and E) Delaunay clustering and nuclear centroid offset was calculated on the cells in the CA1 region alone.

| Morphology Parameter | Description | Interpretation in Microglial Activation States |
|---|---|---|
| Cell Area | The 2D projected area of the microglial cell (soma plus processes) in tissue sections. | Larger cell area indicates a cell with extensive processes covering a big territory (characteristic of hyper-ramified microglia). Smaller area is seen in retracted, activated cells; bushy or amoeboid microglia have contracted arbors and thus lower area. |
| Cell Diameter | The cell's maximum span (e.g. Feret diameter) – an approximate "length" across the cell's extent. | A greater cell diameter usually accompanies larger, ramified cells with long processes. Ramified and hyper-ramified microglia tend to have a larger span due to extended processes. In activated morphologies (bushy/amoeboid), the overall cell span contracts as processes retract, reducing the effective diameter. |
| Convex Hull Area | Area of the convex polygon enclosing the cell (all outermost process tips). | Increases with branching extent. Ramified microglia with long outstretched processes have a much larger convex hull area. A bushy or amoeboid cell occupies a smaller convex hull (processes don't extend far), often nearly equal to its cell area. |
| Circularity | A shape roundness metric ($4\pi \cdot \text{Area}/\text{Perimeter}^2$); 1.0 = perfect circle, lower values = more elongated or irregular shape. | High circularity denotes a round, compact shape, such as that of amoeboid or highly reactive microglia. Low circularity signifies an irregular, branching shape – ramified and hyper-ramified cells with many processes are far from round (lower circularity). |
| Solidity | The ratio of cell area to its convex hull area (a value $\leq 1$); low solidity means a shape has branches/voids, high solidity means it is filled-in/compact. | Ramified microglia have low solidity because their fine processes occupy only a small fraction of the convex hull (lots of space between branches). Activated/bushy microglia have higher solidity as the cell becomes more compact and fills its convex outline (fewer gaps). An amoeboid microglia (rounded) approaches solidity $\approx 1$ (very compact). |
| Aspect Ratio | Ratio of the shortest to longest dimension of the cell (minor/major axis length). | A measure of elongation or polarization. Hyper-ramified microglia can exhibit a higher aspect ratio if processes extend more in one direction. In contrast, a bushy/amoeboid microglia tends to have aspect ratio ~1 (approximately equal dimensions) since it is |

| | | |
|---|---|---|
| | | more symmetric and less elongated. Elevated aspect ratio may also indicate a cell oriented/polarized in one direction (e.g. migrating or rod-like morphology). |
| Skeleton Length | Total length of all processes, obtained by skeletonizing the microglial arbor (sum of all branch segments in μm). | Reflects overall process length/ramification. Ramified and especially hyper-ramified microglia have high total skeleton length (long, numerous processes). Activated microglia (deramified/bushy) have shortened processes, reducing total length, and amoeboid microglia (no long processes) have minimal skeleton length. |
| Number of Branch Endpoints | Count of terminal process tips (skeleton endpoints) of a microglial cell. | Indicates how many free branch ends the cell has. Highly ramified microglia have many branch endpoints (multiple slender process tips). This number drops as microglia activate: bushy microglia have fewer, thickened stubby processes, and an amoeboid microglia may have zero or very few endpoints (having retracted almost all processes). |
| Critical Value | The maximum number of intersections made by the microglial branches with concentric circles (Sholl analysis). This peak intersection count quantifies maximal branching density around the cell. | A higher critical value signifies greater branching complexity, typical of ramified microglia with dense arbors. Fewer peak intersections indicate simpler morphology – activated microglia have a lower peak count. |
| Critical Radius | The radius (distance from soma center) at which the Sholl intersection count is maximal (i.e. where branch density peaks). | Extent of arbor spread. A large critical radius means the cell's branches extend far out before tapering, seen in hyper-ramified or extensively branched cells (peak branching occurs farther from the soma). A small critical radius occurs in bushy/activated microglia with short processes because their maximum branch intersection is close to the soma. Thus, activated microglia have a reduced critical radius, while surveilling ramified cells have a larger one. |

| | | |
|---|---|---|
| Branching Index | A quantitative index of branching complexity, based in the increasing number of branches with greater distance from the soma centroid. | Higher branching index values indicate a richly ramified cell where branches extensively arborize beyond the primary trunks, characteristic of ramified microglia. Low branching index denotes fewer secondary branches relative to primary processes seen in activated microglia that have complex somas but occupy less territory with branches. In other words, as microglia retract processes (toward bushy/amoeboid forms), the branching index drops. By contrast, hyper-ramified microglia show elevated branching index (abundant branching relative to their primary processes). |
| Schoenen Ramification Index (SRI) | Another branching complexity metric defined here as the critical value (peak Sholl intersections) divided by the number of branches. | Like the branching index, SRI increases with greater arbor complexity. Ramified microglia have a high SRI, indicating many branch intersections per initial branch. Amoeboid/activated microglia have SRI near zero or low, as there are few branches and low intersection counts. |
| Branch Thickness | Average thickness of processes, calculated as the cell's cytoplasmic area (cell area minus nucleus area) divided by total process length. This gauges caliber of branches. | Thin, filamentary branches (low branch thickness) are typical of surveillant ramified microglia. An increase in branch thickness occurs as microglia become reactive: bushy microglia have shorter but thicker processes. Accordingly, microglia in a hyper-ramified state often still have relatively thin processes, whereas reactive/amoeboid microglia display stouter processes (high branch thickness). |
| Centroid Offset | Distance between the cell's overall centroid and the nucleus centroid (nuclear offset). Reflects how centrally the nucleus is located within the cell. | In a quiescent ramified microglia, the nucleus usually lies near the center of a symmetrically branched cell (small offset). A larger centroid offset indicates the nucleus is off-center, suggesting the cell is polarized or extending more branch mass to one side. This can occur in hyper-ramified states (asymmetric branch growth) or when microglia polarize towards a stimulus/injury. |
| Iba1 Intensity (Mean) | Average fluorescence intensity of Iba1 (a microglial marker) within the cell. Higher values indicate more Iba1 | Iba1 is generally upregulated with microglial activation. Amoeboid/bushy active microglia often show higher Iba1 intensity (reflecting cytoskeletal activation and cell enlargement). |

| | | |
|---|---|---|
| | protein. | Ramified microglia in resting state have lower Iba1 intensity. |
| CD68 Intensity (% Area) | Lysosomal marker. CD68 measured as % area of the cell that is CD68-positive. High values mean the cell has more CD68+ lysosomal content. | CD68 is low in surveillant/resting microglia and rises as microglia become phagocytic. Reactive microglia (bushy/amoeboid) exhibit high CD68 (intense lysosomal labeling), reflecting active phagolysosomal activity. Homestatic microglia have minimal CD68. |
| Microglial Density | The number of microglia per unit area of tissue (cells/mm²). | Higher microglial density can indicate a proliferative or inflammatory response (microgliosis). An increased density often accompanies activation in disease models, sometimes with clustering. |
| Nearest-Neighbor Distance (NND) | The average distance from one microglia to its nearest microglial neighbor, to the nearest microglial cell, computed via Delaunay triangulation. | NND is inversely related to density: a high average NND means microglia are far apart (low density), while a low NND means cells are closer together (high density). Activated microglia often cluster around lesions, which would reduce NND in those regions. By contrast, uniform surveillant microglia maintain more regular spacing. NND helps distinguish if increased cell count leads to crowding or if cells are still evenly spaced. |

**Supplementary Table 1**

Summary of parameters used to assess microglial morphology, intensity, and spatial distribution.
